# Suitable seasons: Global monthly habitat suitability for the arbovirus vectors *Aedes aegypti* and *Aedes* albopictus in 1975–2024

**DOI:** 10.64898/2026.04.17.719149

**Authors:** Tarique Siddiqui, Nadezhda Malysheva, Anna-Maria Hartner, Semyon Butyrin, Diogo Parreira, Jakob-Wendelin Genger, Christopher Irrgang

## Abstract

The mosquito species *Aedes aegypti* and *Aedes albopictus* are the primary vectors of the arboviruses dengue, Zika, and chikungunya. Expansion of these vectors into previously non-endemic regions due to climate and environmental changes has accelerated global burden from arboviral diseases.

To combat this, predictive models accurately mapping *Aedes* habitats are essential for epidemiological modelling, effective vector control, and disease prevention. We introduce the Climademic Suitability Model, a machine learning model that delivers monthly global predictions of *Aedes* habitat suitability at 0.25° spatial resolution between 1975—2024. The model leverages integrated climate, land use, human population, and mosquito surveillance data to provide an explainable view of the factors governing habitat dynamics.

SHAP-based explainability analysis identified temperature and dew point temperature as dominant features driving habitat suitability. Long-term analysis reveals a complex global redistribution of expanding and contracting vector habitats. Suitable areas for both species now encompass regions home to over 5 billion people, coinciding with the world’s most pronounced population growth and surpassing projections previously placing this threshold at 2050.

The Climademic Suitability Model serves as a framework for near-real-time vector surveillance, climate scenario projection, and integration into transmission models to advance epidemic preparedness in an era of accelerating environmental change.

## 1 Introduction

The global spread of *Aedes aegypti* and *Aedes albopictus* mosquitoes is becoming an escalating public health challenge. Both species transmit major arthropod-borne viruses (arboviruses), including dengue, Zika, and chikungunya, resulting in millions of symptomatic cases and over 10,000 deaths every year [1–5]. Originally endemic to tropical and subtropical regions, the distribution of these mosquito species has broadened substantially in recent decades, mostly supported by intensified global travel and trade [6–10]. As a result, *Aedes* mosquitoes are increasingly reported in regions where they were previously absent. However, despite growing efforts in mosquito surveillance [11, 12] and the increasing use of citizen-science initiatives [13, 14], the true patterns of mosquito range turnover and, in particular, large-scale seasonal activity shifts may still be overlooked as a result of incomplete sampling and limited surveillance coverage.

Although human-mediated transport drives the introduction of these species into new regions, their establishment, persistence, and seasonal activity are shaped by the interplay of climatic, environmental, and socio-economic factors [15–18].

Previous modelling studies [19] have employed a range of approaches to model habitat of these *Aedes* vectors, including machine-learning methods [20–23] and mechanistic models [24–31] that incorporate climatic and non-climatic covariates across both global and local scales. These efforts produced high-resolution habitat-suitability maps for *Ae. aegypti* and *Ae. albopictus*. However, until recently, such models have largely been constrained to annual temporal resolutions due to the limited availability of temporally resolved mosquito-observation data. Over the past decade, increasing access to weekly and daily datasets has enabled finer-scale temporal modelling, improving predictive assessments of mosquito range and dynamics [32, 33].

Most machine learning approaches require balanced presence–absence datasets. Since mosquito observation data typically consist solely of presence data, many studies have resorted to using artificially generated pseudo-absence data, selecting absence locations based on temperature suitability maps for selected species. Other studies rely exclusively on surveillance programs, which generate explicit presence–absence datasets. These approaches, while useful, can be problematic for invasive species subject to underreporting and incomplete sampling. Additionally, crucial habitat suitability dynamics and driver, such as local seasonal cycles and interactions as well as long-term shifts in seasonal dynamics have remained hidden so far due to the complex methodic challenges and data limitations.

We present the Climademic Suitability Model, a data-driven, spatio-temporally dynamic habitat suitability model that provides both short- and long-term insights into the global change of *Ae. aegypti* and *Ae. albopictus* habitat conditions. The model integrates a one-class (OC) Support Vector Machine (SVM) approach that relies solely on presence data, enabling accurate predictions of habitat suitability and explainable environmental analytics.

Integrating multiple mosquito-observation datasets with high-resolution climate, land use, and population-density variables, we generate global suitability zones at 0.25° (ca. 31 km) spatial resolution. Integrating recent weekly and daily mosquito observation records within an *incremental learning* framework, we were able to construct global habitat suitability at a monthly resolution between 1975 and 2024.

The model and data can support a range of use cases from infectious disease epidemiology, entomology, and climate impact research. Working towards resilient global public health systems [34–36], the model aids proactive vector-control planning by identifying areas where favourable conditions for *Aedes* establishment are emerging, seasonal, or persistent before colonisation occurs. By linking environmental drivers to habitat suitability with quantifiable explainability, this study contributes to a broader understanding of how climate and land cover change are reshaping the ecological niches of *Aedes* vectors and the future landscape of arbovirus transmission risk.

## 2 Data sets

### 2.1 Mosquito observation data Global compendium of *Ae. aegypti* and *Ae. albopictus* occurrence (1960-2014)

The global compendium database compiled by Kraemer et al. (2015) [37] is currently the largest available standardized dataset that provides 42,067 spatially unique sightings of adults, pupae, larvae and eggs of *Ae. aegypti* and *Ae. albopictus*. This database comprises of occurrence data derived from peer-reviewed literature while also including unpublished data from national and international entomological surveys. It contains geo-positioned records of 19,930 sightings of *Ae. aegypti* and 22,137 sightings of *Ae. albopictus* between 1960 and 2014. The mosquito sighting coordinates in the database are either linked to specific point locations with a horizontal spatial resolution of 5 km x 5 km or to variable polygon locations that range up to 111 km x 111 km.

#### Global Mosquito Observations Dashboard (2015-2023)

Global Mosquito Observations Dashboard (GMOD) [38] is an interactive web interface that provides open access mosquito observation and habitat data. This dashboard integrates mosquito observation data from three citizen science platforms: GLOBE Observer (Mosquito Habitat Mapper and Land Cover) [39], Mosquito Alert [40] and iNaturalist [41] to create a centralized data repository that provides information on locations as well as dates of mosquito sightings. GMOD contains geo-positioned records of ∼ 20,000 sightings of *Ae. albopictus* and ∼ 4,500 sightings of *Ae. aegypti* between 2015 and 2023 and these records are used in this study to augment the mosquito sightings in the global compendium database [37].

#### Mosquito observations dataset (2024)

Additionally, and primarily for validation purposes (Fig. 1), we compiled mosquito observation data for the year 2024 from several publicly available sources. These included the ECDC Geocatalogue [42], the GBIF Occurrence dataset [43] which aggregates records from surveillance programs, scientific publications, and citizen science initiatives, as well as the dataset from Taiwanese mosquito observation program [44], and the recently launched Brazilian mosquito monitoring initiative Conta Ovos [45].

**Fig. 1:**
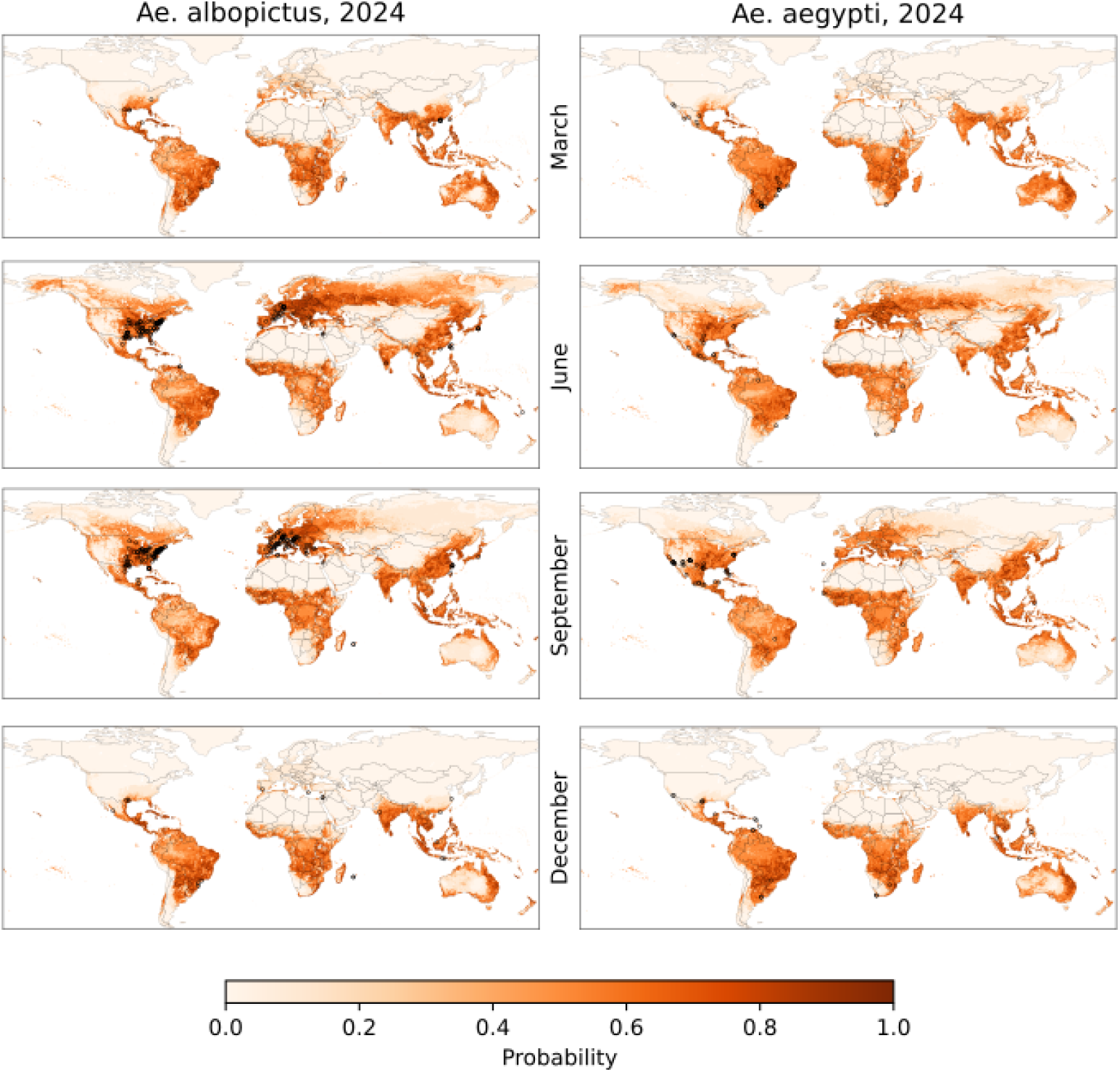
Monthly habitat suitability for *Ae. albopictus* (left) and *Ae. aegypti* (right) in March, June, September, and December of 2024. Maps show posterior probability values of the developed suitability model. Black rings indicate mosquito sightings, data based on validation dataset. Monthly maps for all 12 months of 2024 are shown in SI: Figure 5.

The datasets were provided at heterogeneous spatial resolutions, ranging from precise geographic coordinates to observations aggregated at administrative levels (e.g., municipalities at different hierarchical scales). For this study, observations reported at the highest spatial resolution (exact point locations) were used directly, whereas observations available only at coarser spatial resolutions were represented by the geographic centroids of the corresponding administrative units.

### 2.2 Climate data

ERA5 is the fifth generation global reanalysis dataset produced by European Centre for Medium-Range Weather Forecasts (ECMWF) and currently covers the period from 1940 to the present [46]. ERA5 provides hourly estimates for a large number of global atmosphere, land surface and ocean-wave quantities at a horizontal resolution of 0.25° (ca. 31 km). ERA5 has 137 vertical levels, extending from the Earth’s surface up to 80 km altitude. For this study, we used monthly mean values of surface air temperature (2-m height), dew-point temperature (2-m height), total precipitation and wind speed (10-m height) from the ERA5 dataset. The climatic variables are extracted at coordinates nearest to the central latitude and longitude values of all point and polygon locations of mosquito sightings.

### 2.3 Land use / land cover data

The Historic Land Dynamics Assessment Plus (HILDA+[47]) is a comprehensive dataset that provides annual global land use and land cover data, along with information on changes over time. It is based on a data-driven reconstruction methodology that integrates multiple open data sources, including high-resolution remote sensing data, long-term land use reconstructions, and statistical records. The dataset covers the period from 1960 to 2019, offering detailed information on land use/cover states at a spatial resolution of 0.01° (or 36 arcsec). It classifies land into six generic categories: urban areas, cropland, pasture/rangeland, forest, unmanaged grass/shrubland, and sparse or no vegetation.

### 2.4 Population data

The GHS-POP dataset [48] is part of the Global Human Settlement Layer (GHSL) project and provides high resolution data on the distribution of the global population, expressed as the number of people per grid cell. It includes worldwide population estimates for the years 1975 to 2020 at 5-year intervals, with projections extending to 2025 and 2030. The dataset is available in various spatial resolutions; for our analysis, we chose the 30-arcsecond resolution to ensure compatibility with another dataset and meet the specific requirements of our study. To account for the years between the 5-year intervals, linear interpolation was applied to estimate population counts for each grid cell. The datasets used for model development in this study have been summarized in Table 1.

**Table 1:** Summary of data sets used for model development.

| **Data** | **Source** | **Spatial resolution** | **Period** |
| --- | --- | --- | --- |
| Climate | ECMWF | $0.25^{\circ} \times 0.25^{\circ}$ | 1940 - present |
| Land use / Land cover | HILDA+ | $0.01^{\circ} \times 0.01^{\circ}$ | 1960 - 2019 |
| Population | GHS-POP | $0.083^{\circ} \times 0.083^{\circ}$ | 1975 - 2030 |
| Mosquito sightings | Kraemer et al. (2015) [37] | 5 km × 5 km (points) -<br>111 km × 111 km (polygons) | 1960 - 2014 |
| Mosquito sightings | GMOD | Point locations | 2015 - 2023 |
| Self-compiled<br>mosquito sightings | ECDC, GBIF, local authorities | Point locations | 2024 |

**Table 2:**
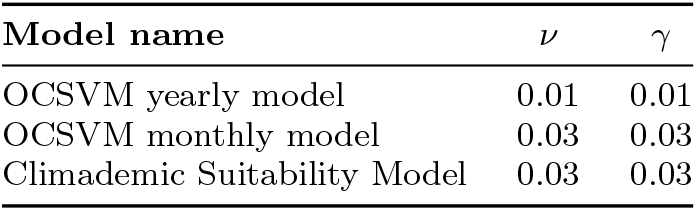
Choice of hyperparameters for OCSVM models.

Due to the compatibility and availability of the data, the analysis in this study covers the period 1975–2024. As land use/land cover data were not available for 2020-2024, it was assumed that the values for these years were identical to those of 2019.

## 3 Methods

### 3.1 One-Class Classification

For binary classification problems with an even distribution of positive and negative classes, traditional machine learning (ML) algorithms aim to build a model that can accurately predict the class of new and unseen instances. However, typical binary classifiers may be unsuitable for classifications tasks when there is class imbalance such that the number of positive instances (class of interest) is disproportionately smaller than the number of negative instances. For such skewed distributions, binary classification algorithms become biased towards the majority class, start to treat the minority class as noise and fail to learn the patterns associated with it. Further, if the availability of data is only for one class and the data of other class is unavailable or unlabelled then for such scenarios traditional binary classification techniques are not the preferred choice of algorithms, rather approaches based on one-class classification (OCC) are the recommended option [49].

OCC is a specific type of classification problem where the goal is to identify objects of a specific class, usually the class of interest, by learning from a training set that contains only examples of that class. OCC is primarily used for anomaly or outlier detection as the models created using this approach effectively learns what the target class looks like and are able to detect instances that deviate from this norm. The One-Class Support Vector Machine (OCSVM)[50], an unsupervised learning algorithm, is one of the most widely used OCC algorithms. OCSVM is a special case of the conventional two-class SVM algorithm, which is based on the idea of finding an optimized decision boundary (hyperplane) that maximizes the margin of separation between the two classes for classification tasks. The creation of the optimal hyperplane is influenced by only a small subset of training samples known as support vectors (SVs), which are the data points from each class nearest to the hyperplane. SVMs were initially introduced for linearly separable problems but have also been extended to handle non-linearly separable problems through the use of kernels, which are mathematical functions that transform the data from a given space (known as input space) to a new high dimensional space (known as feature space) where data can then be linearly separated.

OCSVM algorithm can be formulated mathematically similar to the two-class SVM by treating the training points as one class of data and the origin as the only example of the data of the other class [51]. The algorithm aims to iteratively find the maximal margin hyperplane that best separates the training data from the origin. It focuses on learning the characteristics of only one-class of data for training with the goal of detecting whether a new data instance differs significantly from the distribution of the training data.

### 3.2 OCSVM yearly dataset preparation and model development

For the development and training of the OCSVM yearly model, we create the `Climate_LULC_Population_Yearly` dataset by selecting climate, LULC and population variables corresponding to each instance of yearly mosquito sighting. Due to the different temporal coverage periods of mosquito occurrence, climate, LULC and population datasets, we choose the years common to all of them, i.e. between 1975 and 2014, for our analysis. As a result, the number of mosquito sightings reduces to 18,518 for *Ae. aegypti* and 21,703 for *Ae. albopictus* for the 1975-2014 analysis period.

While LULC and population data are themselves available at yearly resolutions, climate variables are available at monthly resolution and are chosen as follows: monthly means of temperature, dew-point temperature, total precipitation and wind speed are extracted for the months of January to December for each instance of yearly mosquito sighting. By selecting monthly means from January to December, we ensure that every month of the year is equally represented in the four climate variables for every instance of mosquito sighting. In total, we combine 48 (12 x 4) climate variables, 8 LULC variables and 1 population variable to create 57 (48+8+1) features for the occurrence record of mosquito sightings, which is the target variable.

The OCSVM yearly model is developed using the `Climate_LULC_Population_Yearly` dataset after applying the data splitting and standardization preprocessing steps. The prepared dataset is partitioned using a 80/20 split with a fixed random state in order to obtain training and test sets. The features in the training and test sets are standardized using the Z-score method (Standard Scaler from the Scikit-learn library). The scaler is first fitted only to the training data using the fit transform method and then the test data is transformed using the parameters learned from the training data to avoid any data leakage from the test set. The OCSVM yearly model is configured with a radial basis function (RBF) kernel as it is well-suited for capturing non-linear relationships in datasets and is the most versatile go-to kernel in SVM because it can model a wide range of complex data distributions [51]. The OCSVM model also requires two hyperparameters, gamma (*γ*) and nu (*ν*), to be properly tuned as they have significant influence on its performance. The kernel coefficient parameter, *γ*, influences the shape of the decision boundary of the model with a smaller *γ* creating a wider, smoother and more generalized boundary, while a larger *γ* resulting in a tighter, more specific and complex decision boundary. The contamination parameter, *ν* ranges between 0 and 1 and controls the trade-off between the number of support vectors and the number of training errors. It determines the lower bound on the fraction of SVs and the upper bound on the fraction of data points that the OCSVM model should treat as outliers. The tuning of OCSVM hyperparameters is not straightforward since standard hyperparameter selection methods such as cross-validation cannot be directly applied to OCC methods due to the absence of data from outlier class. OCSVM hyperparameter selection in past studies have usually relied on heuristic based methods that typically satisfy some prior empirical knowledge. We employ a heuristic estimate by assuming that only 1% of the training data can be treated as outliers. Based on this assumption, we set the value of *ν* to 0.01. The value of *γ* is determined using typical data-driven approaches such as the median heuristic technique, which is defined as the inverse of the median of all pairwise distances between data instances [52–54]. The median heuristic technique is a common approach for determining the *γ* parameter of the RBF kernel [54]. The value of *γ* is estimated to be 0.01 with this method and the accuracy over training data reaches the desired 99% score for the choices of *γ* = 0.01 and *ν* = 0.01. The performance of the OCSVM yearly model cannot be evaluated using classical criteria such as Receiver Operating Characteristic (ROC) or Precision-Recall (PR) curves since these techniques only work when sufficient labelled data are available [55]. In the case of OCSVM yearly model, the test set is unlabelled and contains data of only one class, which makes it impossible to create ROC curves. The evaluation of the OCSVM yearly model is therefore done based on the stability criteria by checking if the model produces similar score distributions on the test and training data sets. The OCSVM yearly model is found to fulfil the stability criteria as it also returns a 99% accuracy score for the choice of hyperparameters, *γ* = 0.01 and *ν* = 0.01, on the test data set and therefore represents a generalized model. With these empirical estimations of hyperparameters, the OCSVM yearly model has been developed in this study.

For the validation of OCSVM yearly model, we first create yearly datasets beginning from the year 2015 by concatenating the climate, LULC and population variables over all global coordinates at a horizontal resolution of 0.25° × 0.25°. Similar to the training data set, the validation data sets contain 48 (12 x 4) climate variables, 8 LULC variables and 1 population variable. The availability of the HILDA+ dataset limits the validation period to 2019. However, with the assumption that LULC variables do not change substantially over a 5 year period, the validation period has been extended until 2024. Hence, for the years 2020-2024 we concatenate the respective climate and population variables but keep the LULC variables to the levels seen during the year 2019.

The output of a OCSVM model for each data point is a decision score whose value quantifies whether an instance is an inlier or an outlier. For a data point, the decision score represents its distance to the learned decision boundary in the feature space. A positive value signifies that the data point is an inlier while a negative value suggests that the data point is an outlier and falls outside the region learned by the model. For the validation datasets, created for the years 2015-2024, we use the OCSVM yearly model to obtain the classification of global coordinates into inliers and outliers at a horizontal resolution of 0.25° × 0.25°. The global maps showing the coordinates that are inliers and outliers are plotted for the validation datasets for the year 2024 for both *Ae. albopictus* and *Ae. aegypti* in SI: Figure 1. The points in red are the ones that are classified as inliers by the OCSVM yearly model while the points in grey are classified as outliers. The inlier points denote the locations that are predicting habitat suitability for *Ae. aegypti* and *Ae. albopictus* while the outlier points denote the locations that are unsuitable for these two species.

The next step after obtaining the binary classification outputs of inliers and outliers from the OCSVM yearly model involves the mapping of OCSVM decision scores to actual probabilities. The OCSVM model does not inherently provide probabilistic outputs but the decision score that it produces for each data instance can be converted to a probability-like output. Earlier works that have used the decision score of data instances to predict the probability have generally been restricted to studies where the classification method was based on two-class SVM [56–58]. Among them, Platt scaling [56] is a popular method that has been frequently used to convert decision scores of two-class SVM into probabilistic outputs. However, obtaining probabilistic outputs from one-class SVM has remained a challenge mainly because of lack of labelled information. It has been recently shown by Que and Lin (2024) [59] that existing methods used to obtain probabilistic outputs for two-class SVM may not be suitable for one-class SVM. Their results further showed that allowing the probabilities to mimic the decision scores of training data is a better approach to generate probabilistic outputs for one-class SVM. The method they propose involves binning the decision scores by dividing the range of decision values into several intervals and then assigning the same probability to the samples with decision scores in the same interval. Further, to offset the effect of outliers, Que and Lin (2024) [59] suggest to generate the binning intervals according to the density of the decision scores. This non-parametric approach of obtaining probability values from OCSVM decision scores has also been implemented in the popular LIBSVM package. With the quantile binning approach employed in LIBSVM, the OCSVM decision scores are partitioned into 10 subsets of approximately equal size. On a horizontal grid resolution of 0.25° × 0.25° employed in this study, the number of counts per bin approximate to roughly 20,000, which results in an uncertainty in the bin occupancy of less than 1% under the assumption of Poisson statistics. We follow the approach of Que and Lin (2024) [59] in this work and use the LIBSVM package to map the decision scores out of OCSVM yearly model to probability values for both *Ae. albopictus* and *Ae. aegypti* and present them in SI: Figure 2.

**Fig. 2:**
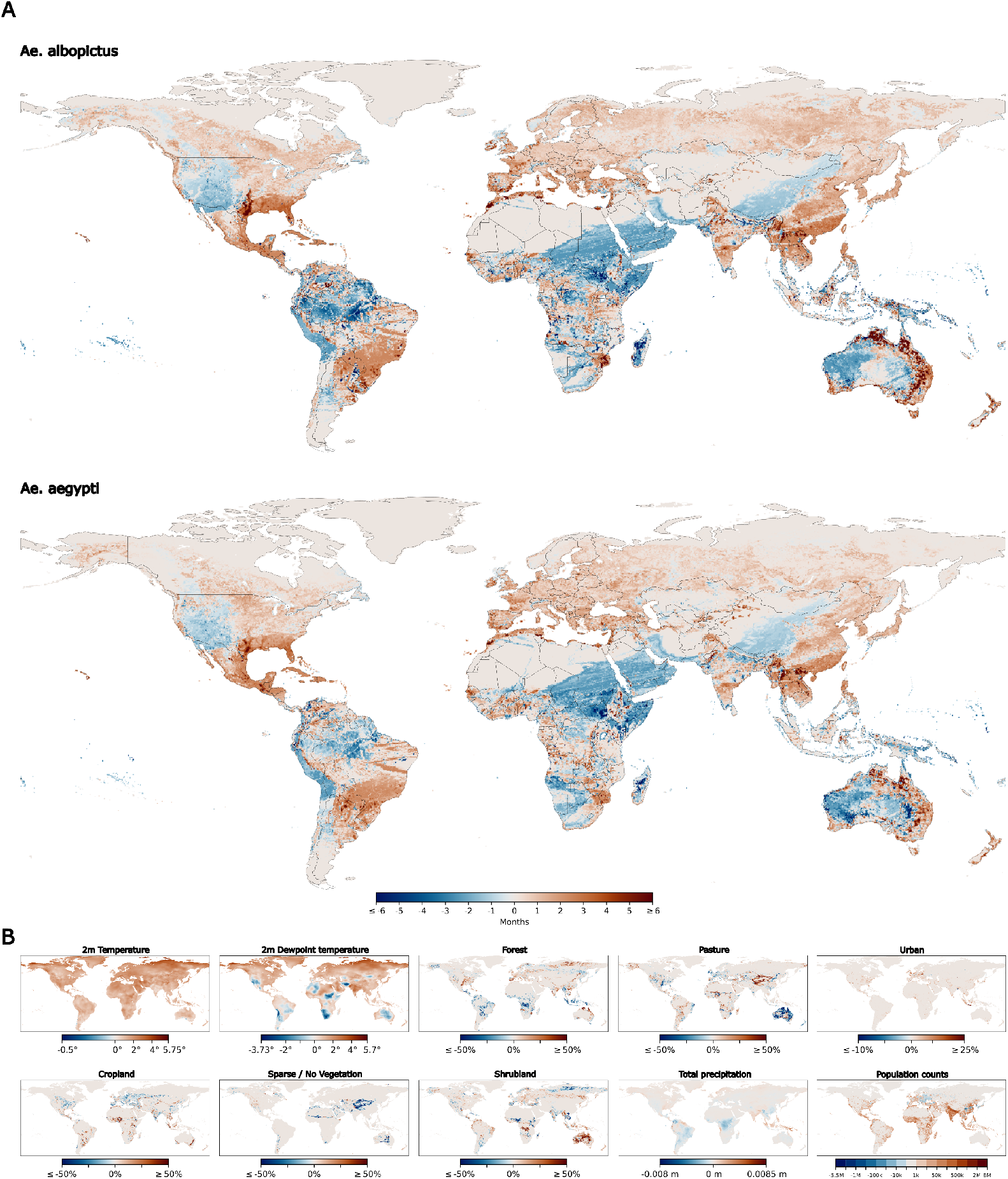
A. Long-term changes in seasonal habitat suitability of *Ae. albopictus* (top) and *Ae. aegypti* (bottom). The maps show habitat suitability shifts in months for the decade 2015–2024 compared to the decade 1975–1984. Positive (negative) values indicate regions of increasing (decreasing) seasonal habitat suitability over the last 50 years. B. The maps show long-term changes over the same two time periods for the suitability model features.

### 3.3 OCSVM monthly dataset preparation and model development

The `Climate_LULC_Population_Monthly` dataset is prepared in a manner similar to its yearly counterpart but contains comparatively fewer features. Corresponding to each instance of mosquito sightings, LULC and population density data are selected for the monthly dataset in an identical way as the yearly one, but the selection of climate variables is different from the yearly one both in terms of their number and the manner in which they are chosen in the monthly dataset. Unlike the yearly dataset, where all twelve monthly mean values of climate variables are selected as features for mosquito occurrence records, we assign single attributes of the four climate variables as features in the monthly dataset. This approach allows us to eventually input only one value for climate variables, i.e., monthly mean values of temperature, dew-point temperature, total precipitation and wind speed to generate corresponding monthly prediction of habitat suitability for *Ae. aegypti* and *Ae. albopictus*. However, choosing single attributes of the four climate variables that can be related to individual yearly mosquito sightings is not straightforward as the specific month in which the individual mosquito sightings were recorded remain unknown and are not mentioned in the global compendium database. We employ a heuristic approach to hypothesize an approximate estimate of the seasonal time frame and the climate conditions in which the individual mosquito sightings were recorded. Corresponding to the year of each mosquito sighting, monthly means of temperature, dew-point temperature, total precipitation and wind speed are extracted for the months of January to December and the third quartile (75th percentile) value of these twelve monthly means are calculated for each of the four climate variables. The 75th percentile value of climatic variables has been chosen in order to capture the optimal and conducive climatic conditions for *Aedes* mosquitoes that are relatively warmer and wetter than the median conditions and therefore can justify a reasonable initial estimate of the approximate climatic conditions for mosquito sightings. We combine the values of the 4 climate variables with the values of 8 LULC variables and 1 population variable to create the `Climate_LULC_Population_Monthly` dataset that contains 13 (8+1+4) features for the target variable, which is the occurrence record of mosquito sightings.

Similar to the development of OCSVM yearly model, the development of OCSVM monthly model involves partitioning the `Climate_LULC_Population_Monthly` dataset into training and test set using a 80/20 split with a fixed random state. Thereafter, the features in the training and test sets are standardized using the Z-score method (Standard Scaler from the Scikit-learn library). The scaler is first fitted only to the training data using the fit transform method and then the test data is transformed using the parameters learned from the training data. Like the OCSVM yearly model, the OCSVM monthly model is configured with a radial basis function (RBF) kernel. For the monthly data set, we heuristically assume that 3% of the training data can be treated as outliers. Based on this assumption, we set the value of *ν* to 0.03. The value of *γ* is determined using the median heuristic technique and its value is found to be 0.03. The accuracy over training data reaches the desired 97% score for the choices of *γ* = 0.03 and *ν* = 0.03 and the OCSVM monthly model also returns 97% accuracy score on the test data set. Therefore, with these empirical estimations of hyperparameters, the OCSVM monthly model has been developed in this study.

The decision of choosing 75th percentile value of climatic variables and the above mentioned values of hyperparameters is based on the prior knowledge about the known effects of environmental temperature on the fundamental metabolic and ecological processes of both *Aedes* mosquitoes. Environmental temperature is the primary climatic factor shaping the ecology of *Ae. aegypti* and *Ae. albopictus*. Their survival, feeding behaviour, and development are all strongly temperature-dependent.([60, 61]. Past reports have shown that the survivability limits of both *Ae. aegypti Ae. albopictus* generally do not exceed beyond the temperature ranges between 10 °*C* and 36 ° *C* (e.g., [61],[62],[63],[64]). In SI: Figure 3, it is shown that the support vectors satisfy the temperature survivability limits of both *Ae. aegypti Ae. albopictus* with the choice of 75th percentile value of climatic variables and the values set for hyperparameters. Instead of 75th percentile value of climatic variables, if the 50th percentile value is chosen then the support vectors go beyond the temperature survivability limits of both *Ae. aegypti Ae. albopictus* as shown in SI: Figure 4.

**Fig. 3:**
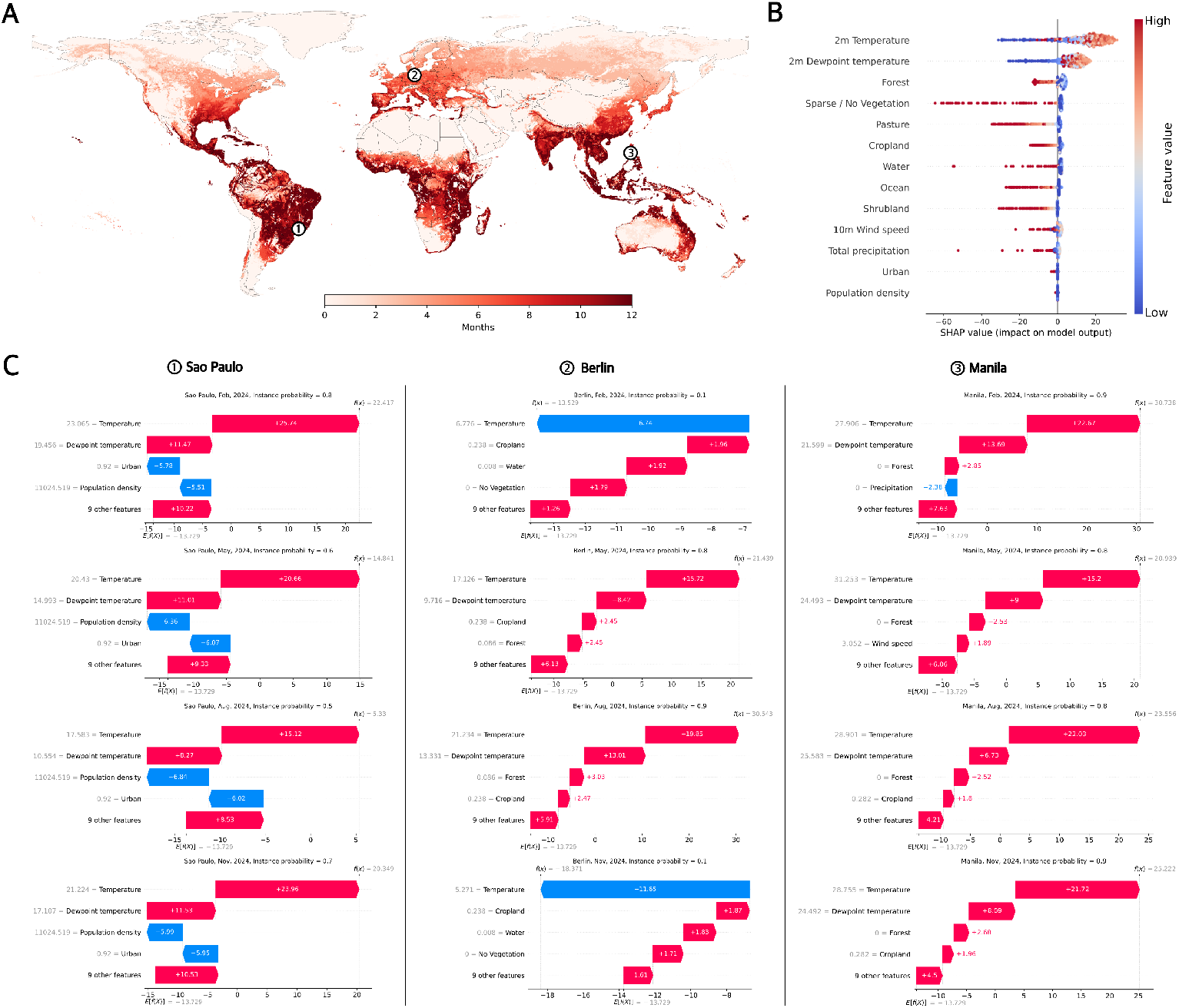
Habitat suitability explainability analysis for the case of *Ae. albopictus* in the year 2024. A. Worldwide display of monthly habitat suitability in 2024. B. SHAP (SHapley Additive exPlanations) impact analysis of the most important model features for the year 2024. C. Location-specific SHAP waterfall plots for the locations of Sao Paulo, Berlin, and Manila (see columns and map indicators 1 to 3), each shown for the months of February, May, August, and November of 2024 (top to bottom).

**Fig. 4:**
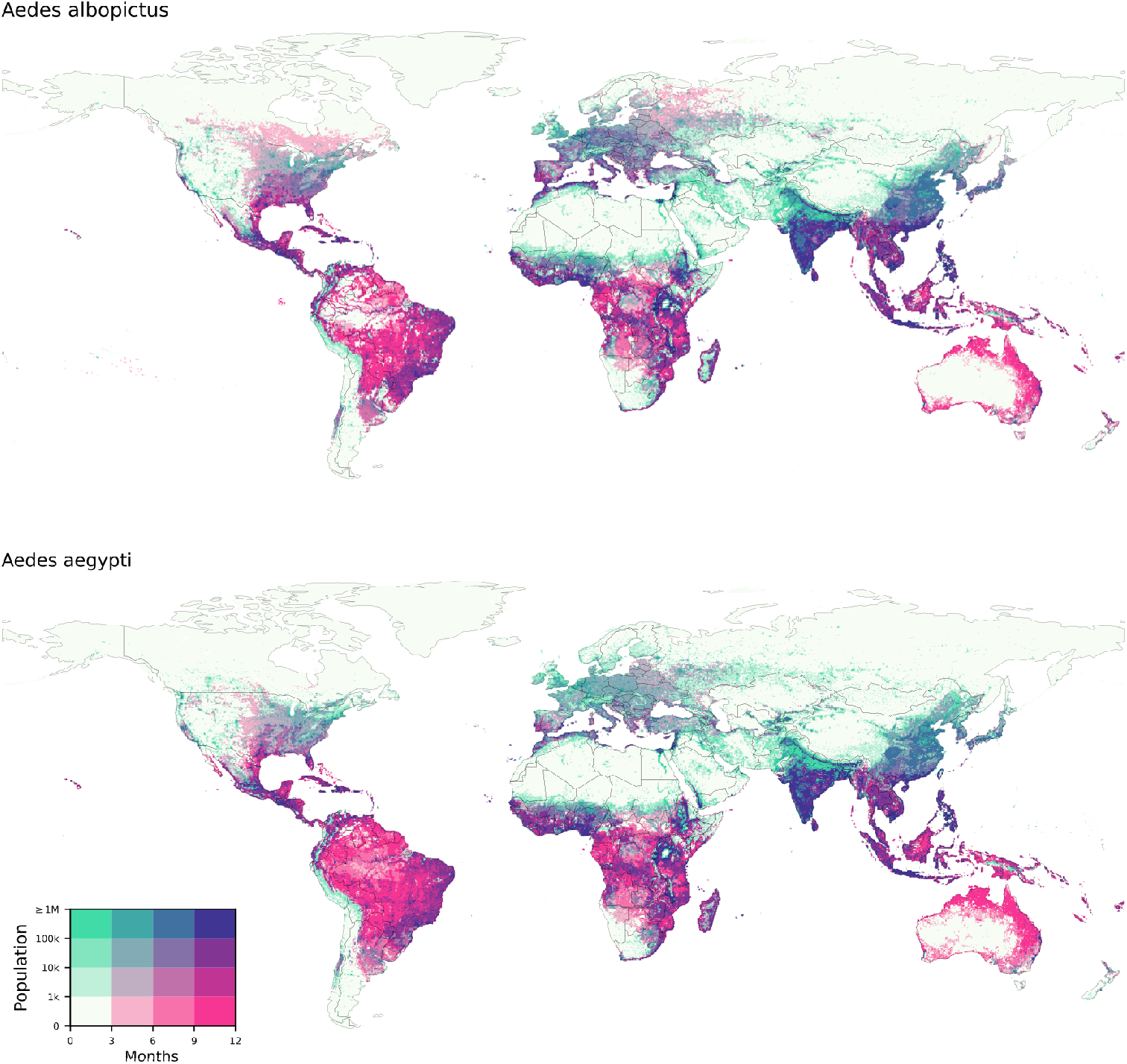
Bivariate choropleth plot of binned global population and binned monthly habitat suitability for *Ae. albopictus* (top) and *Ae. aegypti* (bottom) in 2024. Light colors (lower left part of the color matrix) indicate less populated regions with shorter habitat suitability, dark regions (upper right part of the color matrix) indicate highly populated regions with extended habitat suitability.

The OCSVM monthly model is initially used to obtain global maps that show the classification of data points into inliers and outliers at a horizontal resolution of 0.25° × 0.25° and later these maps are converted into probabilities by binning the OCSVM decision scores in a similar way as described earlier in more detail for the OCSVM yearly model. Figure 5 shows the monthly global habitat suitability maps for all the twelve months of 2024 for *Ae. aegypti* and *Ae. albopictus*, respectively.

### 3.4 Climademic Suitability Model: Integration of GMOD dataset with the OCSVM monthly model

The availability of mosquito sightings between 2015 and 2023 through GMOD provides a new data set that can be possibly used to refine the existing OCSVM monthly model. We aim to sequentially integrate these new observations from GMOD to refine the existing decision boundary of the OCSVM monthly model. To achieve this goal, the same set of climate, LULC and population features that are included in the `Climate_LULC_Population_Monthly` dataset are first extracted for each new mosquito sighting from GMOD. As the GMOD dataset provide the dates of mosquito sightings, the specific monthly values of climate variables that correspond to these observations are obtained and joined with the yearly values of LULC and population variables for each instances. As mentioned earlier during the creation of the `Climate_LULC_Population_Monthly` dataset, the LULC and population variables are assumed to be constant throughout the year and therefore their yearly values are extracted for the mosquito sightings in GMOD.

The GMOD data set is sequentially integrated into the OCSVM monthly model using an incremental learning approach. Incremental SVM techniques have been employed in prior studies to overcome computational challenges associated with either extremely large training datasets or the continuous, staggered arrival of new data [65, 66]. The incremental learning approach is applied in an iterative manner in order to improve the decision boundary of the OCSVM monthly model trained using 75th percentile value of climatic covariates associated with the mosquito sightings[37]. As the training data of the OCSVM monthly model span from 1975 to 2014, the GMOD data for the year 2015 is first fed into the OCSVM monthly model (hereafter referred to as OCSVM_*t*_) for classification and only the misclassified data are retained for incremental learning while the correctly classified data are discarded. Thereafter, the support vectors of OCSVM_*t*_, together with the misclassified GMOD data for the year 2015, are used as training data to obtain a new model OCSVM_*t*+1_, which is configured with the same hyperparameter values of *ν* and *σ* as OCSVM_*t*_. This method was subsequently applied iteratively to the GMOD data spanning the period from 2016 to 2023 and a new updated model was obtained yearly. The final OCSVM monthly model obtained after the ninth iteration is coined as the Climademic Suitability Model. This approach is also beneficial as it creates a pipeline for further annual updates in order to investigate the yearly differences in habitat suitability.

The Climademic Suitability Model is evaluated on its ability to correctly classify the habitat suitability of *Ae. albopictus* and *Ae. aegypti* by comparing the predicted probabilities for the locations of these mosquitoes sightings for the year 2024. Probability levels of 0.5 correspond to points on the decision boundary and points that are outside this decision boundary have probability values less than 0.5. The performance of the model is evaluated with two criteria: classification accuracy and mean absolute error. The classification is considered to be accurate if the locations of mosquito sightings are classified as inliers by the model. For the calculation of mean absolute error, however, the actual probability values predicted by the model are used and evaluated against the probability values of 1 that are assigned for mosquito sightings. The performance of the model for *Ae. albopictus* and *Ae. aegypti* sightings for the year 2024 are summarized in Table 3.

**Table 3:** Performance evaluation of Climademic Suitability Model for the year 2024.

| Specie | Number of sightings | Accuracy | MAE |
| --- | --- | --- | --- |
| *Ae. albopictus* | 2534 | 0.91 | 0.27 |
| *Ae. aegypti* | 2775 | 0.95 | 0.21 |

### 3.5 Model explainability

We utilize SHapley Additive exPlanations (SHAP) [67] method to understand the outputs and to quantify the importance of individual features in the Climademic Suitability Model. The concept of Shapley values has been originally adapted from cooperative game theory and is used to determine the fair contribution of each player to the total outcome of a collaborative game [68]. In the context of machine learning, this concept is utilized to explain the prediction of a model by analogically treating the features as the players and the value of prediction as the total outcome. The calculation of Shapley values rely on the machine learning model trained on the given dataset. The Shapley value of a feature is calculated as its mean marginal contribution to all possible coalitions. For *n* features, this results in 2^*n*^ coalitions and this creates a major challenge because the calculation of exact Shapley values becomes computationally challenging for data sets with large number of features [69].

The recommended algorithmic-specific SHAP variant to use for OCSVM is KernelSHAP, which is a model-agnostic technique to explain the predictions especially when a non-linear kernel such as RBF is used [67]. KernelSHAP provides an approximation of the exact Shapley values by sampling feature coalitions based on a background data distribution instead of evaluating all possible coalition subsets [67]. The background data distribution is used by KernelSHAP to simulate the exclusion of features in the coalitions as well as to derive the base expected value of the model, which is the average prediction of the model over the background dataset. The base expected value serves as the baseline to which the Shapley values of all *n* features are summed for explaining the model output of a specific instance.

The visualization tool in the SHAP framework provides both global and local interpretation capabilities. Global interpretation can be visualized using feature importance and summary plots while local interpretation can be done through waterfall plots to gain an understanding about the feature contributions for a single observation through individual Shapley values. We utilize these tools to explain and interpret the results of the Climademic Suitability Model.

## 4 Results

### 4.1 Model predictions

#### 4.1.1 Seasonal variability

The Climademic Suitability Model is utilized to create global habitat suitability maps for *Ae. albopictus* and *Ae. aegypti* for the validation year 2024. Figure 1 shows these maps for the months of March, June, September and December for *Ae. albopictus* (left side) and *Ae. aegypti* (right side). These months have been chosen in order to assess the capability of the model to capture the seasonal variability that is associated with the habitat suitability of *Ae. albopictus* and *Ae. aegypti*. In the Northern Hemisphere, with the progression of the season from boreal spring (March) to boreal summer (June), the distributions of both species over the Northern Hemisphere show an increase in habitat suitability. The habitat suitability maps extend farther into the northern latitudes in June than in March for both *Ae. albopictus* and *Ae. aegypti*. As the season in the Northern Hemisphere changes to boreal autumn (September), the latitudinal extent of the habitat suitability of both species decline in comparison to June and get further diminished in boreal winter (December). In contrast to the Northern Hemisphere, the model predictions show that the latitudinal extent of the distributions of the two species in the Southern Hemisphere first decrease as the season changes from austral autumn (March) to austral winter (June) and then progressively increase through austral spring (September) and austral summer (December). The habitat suitability maps predict the distributions of the two species to have similar expanse over much of the global regions during these four months. However, there are some notable differences in parts of Europe, North America, Brazil, Madagascar and Australia as *Ae. albopictus* and *Ae. aegypti* exhibit distinct distributions in these regions. Over Europe and North America, *Ae. albopictus* seems to demonstrate greater habitat suitability than *Ae. aegypti* while in Brazil, Madagascar and Australia, *Ae. aegypti* is found to have geographically widespread habitat than *Ae. albopictus*.

The model predictions across March, June, September and December are compared with the actual sightings of *Ae. albopictus* and *Ae. aegypti* for the year 2024, which are shown using black circles in Figure 1. In a qualitative sense, the model results capture the extent of the actual mosquito sightings of these *Aedes* mosquitoes with the predictions being particularly accurate over North America for both *Ae. Albopictus* and *Ae. aegypti* for the months of March and December. For these months, the model results are able to reliably capture the geographical extent of the mosquito sightings over North America for both *Aedes* species. The model results also correctly capture the geographical extent of *Ae. aegypti* over Brazil and South Africa for the months of March, June and December. Over Europe and North America, the model predictions, in general, overestimate the actual mosquito sightings for the months of June and September. Although the model predicts the potential distribution of *Ae. albopictus* over Canada, Russia, Scandinavia and Balkans, the presence of this specie is yet to be reported in these regions. Similarly the distribution of *Ae. aegypti* over Europe as predicted by the model has not yet been validated by sightings. The results of these maps should be considered based on the context that many of the regions that support the habitat suitability of these *Aedes* species remain yet to be colonized. For specific months, the habitat suitability maps of these *Aedes* species broadly inform about the fact that these species would be able to survive if they were introduced in these areas in these particular months.

#### 4.1.2 Long-term changes in habitat suitability

The Climademic Suitability Model is further used to analyse the global evolution and long-term changes in the habitat suitability of *Ae. albopictus* and *Ae. aegypti* over the past decades. For this purpose, we compute monthly model predictions for time intervals 1975–1984 and 2015–2024.

The global maps showing these long-term decadal changes for *Ae. albopictus* and *Ae. aegypti* are shown in Figure 2A. Broadly, for both *Ae. albopictus* and *Ae. aegypti*, the areas of the world that show a net positive or negative change in the average number of suitable months are mostly common. From the figure, the global trend of expanding habitat suitability of *Aedes* mosquitoes can be mainly seen in central America, eastern United States, Europe, southern Brazil, parts of India, eastern China, south-east Asia, Japan and eastern Australia. The noticeable decrease in habitat suitability is visible in western United States, northern Brazil, large parts of Africa, central China and western Australia.

Over the course of past four decades, the model predictions reveal that most of the Mediterranean Europe has seen an average increase in the habitat suitability of both *Aedes* mosquitoes by at least 2-3 months while over central and eastern Europe, the increase in habitat suitability is by 1-2 months. Over sub-saharan Africa and Saudi Arabia, the model results predict a strong average decline in habitat suitability by more than 3 months. In southern Brazil, the model results reveal an average increasing suitability of both *Aedes* mosquitoes by 2-3 months while a similar decrease is seen in northern Brazil. The increase in suitability is also seen in central America and south eastern United States by at least 3 months while in western United States, there is a decline by up to 2 months. Over south-east Asia and eastern China, there is an increase in habitat suitability by up to 3 months while there is a decline in habitat suitability in western China. In Australia, there is a noticeable increase in habitat suitability of both *Aedes* mosquitoes in the east coast and an equally strong decline on the west coast.

To explore the covariates that could qualitatively explain the long-term decadal changes in the habitat suitability of *Ae. albopictus* and *Ae. aegypti*, we compute the average change in the features between the 2015–2024 and 1975–1984 decades and plot it in Figure 2B. At a first glance, it is difficult to find any common pattern between the decadal changes in temperature and the decadal changes in habitat suitability as temperature has mostly increased all over the globe in the past four decades as seen in Figure 2B. However, the decadal changes in dew-point temperature seem to show a correlation with the decadal changes in habitat suitability of both the *Aedes* mosquitoes. The areas that have seen a reduction in habitat suitability are mostly aligned with the areas that have seen a decline in dew-point temperature over the past four decades. This pattern is evident over western United States, over the western coast of South America, over central Brazil, in southern and central Africa and in Australia. The long-term changes in the pattern of land use land cover variables can also be used to understand the changes in the habitat suitability over certain regions. For example, the increase in habitat suitability in southern part of Africa, in the region shared by South Africa, Mozambique and Eswatini, can be recognized through the change in this region from being a pasture land to a forest. This change in land use is likely a potential contributor for the increased habitat suitability in this region. The changes in land use resulting in increase of forest areas in the past four decades over Europe, North America, south-east Asia and northern Australia also coincides with the increase in habitat suitability over these regions. Overall, the net change in habitat suitability all over the world is not limited to the decadal changes of a single feature but rather rely on a complex interplay of net change among all features.

#### 4.1.3 Aggregation of habitat suitable months

The number of habitat suitable months for the year 2024 are aggregated and plotted for *Ae. albopictus* as shown in Figure 3A. A similar plot for *Ae. aegypti* is included in SI: Figure 6. From Figure 3A, it can be seen that the model predicts year-round habitat suitability of *Ae. albopictus* in most of the regions of Brazil, central America, sub-Saharan Africa, India, south-East Asia and eastern China. Over United States, *Ae. albopictus* has year-round habitat suitability in parts of Texas and Florida, while in Europe, southern Portugal, Spain and coastal regions of Italy are the areas that are predicted to be year-round habitat suitable for *Ae. albopictus*. The habitat suitability of *Ae. albopictus* is predicted to lie between 6 and 8 months for France, Germany and Poland in 2024 while in Russia, the habitat suitability of *Ae. albopictus* is predicted to be around 4 months in 2024. In Nordic countries and Canada, the model predicts the habitat suitability of *Ae. albopictus* to be ≤ 3 months in 2024. In Australia, the habitat suitability of *Ae. albopictus* is predominantly determined to be on the eastern and northern regions with year-round habitat suitability predicted for most of these areas while in New Zealand, *Ae. albopictus* is seen to be mostly confined to northern regions with a varying habitat suitability between 6 and 12 months.

### 4.2 Model interpretation

The SHAP summary plot of the Climademic Suitability Model for *Ae. Albopictus* for the validation year 2024 is shown in Figure 3B. The summary plot showcases the ranking of the features in accordance to their importance in the y-axis with the feature at the top being the most important and at the bottom being the least important. In the x-axis, the Shapley values of the features are depicted. Each dot on a feature’s row represent the Shapley value of that feature for a single instance while the color of each dot correspond to the value of feature for that instance with red and blue depicting high and low feature values, respectively. In total, there are 100 instances (dots) shown in the summary plot, which have been randomly chosen by stratified sampling of ten instances from each of the ten probability bins. The summary plot shows that temperature and dew point temperature are the two most important features of the Climademic Suitability Model with LULC variables such as forest, sparse/no vegetation, pasture, cropland and water also having a noticeable influence on the model output. In general, moderate values of temperature and dew point temperature, in a relative sense, contribute to the largest positive Shapley values, while low values of these two variables result in negative Shapley values. The Shapley values of LULC variables are highly asymmetrical with their higher features values contributing to large negative Shapley values while for their lower feature values, the Shapley values are closer to zero and thus have relatively less positive impact on the model output than temperature and dew-point temperature.

To obtain a deeper understanding of the Climademic Suitability Model, we use the SHAP waterfall plots to visualize local interpretations of the model and to gain an intuition about how certain features are affecting the decision-making process of the model. To highlight this aspect, we first select three geographically diverse cities, namely, Sao Paulo (Brazil), Berlin (Germany), Manila (Philippines) and then create individual SHAP waterfall plots based on the model outputs for the months of February, May, August and November for the year 2024, which are presented in Figure 3C. The SHAP waterfall plots highlight the features that are responsible for driving the model output from the base expected value (*E*[*f* (*X*)]) to the actual instance value (*f* (*x*)). *E*[*f* (*X*)] is the mean prediction value that is predicted only based on the background data distribution, which simulates the exclusion of all features while *f* (*x*) is the instance output value of the prediction with all features. In the plot, the Shapley value contribution of each feature is either positive or negative and the superposition of all Shapley values lead to the instance value, *f* (*x*), starting from the baseline *E*[*f* (*X*)].

For Sao Paulo (1), the Climademic Suitability Model predicts the instance probability for the month of February to be 0.8. The corresponding SHAP waterfall plot row wise lists the highest positive or negative Shapley values of the top 4 features along with their feature values while the summed Shapley values that represent the combination of 9 other features are listed in the fifth row. Starting from the base expected value, *E*[*f* (*X*)] = − 13.729, it is seen that the summed Shapley values of 9 other features equal 10.22 while population density and the urban LULC variables contribute negatively with Shapley values of -5.51 and -5.78, respectively. The largest Shapley values contributions come from dew-point temperature and temperature with values of 11.47 and 25.74, respectively, and the summation of all these five components lead to the instance value *f* (*x*) = 22.417. In a similar way, the SHAP waterfall plots for Sao Paulo for the months of May, August and November can also be interpreted. For the months of May, August and November, the instance probabilities for Sao Paulo predicted by the model are 0.6, 0.5 and 0.7, respectively, while the values of *f* (*x*) for these three months are 14.841, 5.33 and 20.349, respectively. The relationship between instance probabilities and *f* (*x*) values can be understood in terms of OCSVM decision scores. Higher instance probabilities are associated with higher OCSVM decision scores leading to higher *f* (*x*) values and vice-versa. In case of Sao Paulo, *f* (*x*) values are mostly controlled by the Shapley values of temperature, which itself is seen to vary linearly with the feature values of temperature.

For Berlin (2), the Climademic Suitability Model predicts the instance probability for the months of February, May, August and November to be 0.1, 0.8, 0.9 and 0.1, respectively. The higher probability levels as well as higher *f* (*x*) values in May and August for Berlin are largely associated with positive Shapley values of temperature and dew-point temperature while the lower probability and *f* (*x*) levels in February and November are associated with large negative Shapley values of temperature. The low feature values of temperature in February and November makes these monthly instances to be treated as an outlier by the Climademic Suitability Model and is the main reason behind their lower probabilities and lower *f* (*x*) levels for Berlin in February and November. For Manila (3), the probability levels predicted by the Climademic Suitability Model for the months of February, May, August and November are 0.9, 0.8, 0.8 and 0.9, respectively. The similarity in probability levels for Manila across these months lead to similar SHAP waterfall plots with temperature and dew-point temperature having the largest Shapley values. The contributions of LULC variables do not play a major role in determining the instance probabilities in case of Manila for these aforementioned months.

## 5 Discussion

In this study, we combined climate, land use, and human population datasets with mosquito observation data to estimate global seasonal suitability patterns for *Ae. aegypti* and *Ae. albopictus* species, offering a robust framework for understanding temporal and spatial variation in potential vector risk. To leverage the full extent of the collected dataset, we develop models at a global scale, which enables the integration of insights across diverse environmental regions. Starting from the yearly habitat suitability maps of these species at a 0.25°× 0.25° spatial resolution, we employ a novel mechanistic approach to improve habitat suitability maps to a monthly temporal resolution, which are shown in Fig. 1 for both species for the validation year 2024.

Previous studies have utilised a wide range of modelling approaches and diverse covariates to model the distributions of *Ae. albopictus* and *Ae. aegypti* at different spatial scales [70–74]. In particular, the global maps of *Ae. albopictus* and *Ae. Aegypti* produced by [70] and [73] offer a good baseline for making a comparison with our results. Despite the differences in these modelling techniques, the yearly habitat suitability plots of *Ae. albopictus* and *Ae. aegypti* derived from our approach are largely consistent with similar maps produced by [70] and [73]. The distributions of *Ae. albopictus* and *Ae. aegypti* over North America shown in SI: Figure 1 largely match the results of both [70] and [73], while over Europe, the limited habitat suitability beyond northern Italy shown for both *Ae. albopictus* and *Ae. aegypti* in [70] differs with our results as well as with the findings of [73]. The extended northward distributions of *Ae. albopictus* as compared to *Ae. aegypti* over Europe, United States and Japan as seen in SI: Figure 1 suggests its ability to tolerate lower temperatures than *Ae. aegypti*, which is consistent with the findings of [73].

Using spatio-temporal modelling methods, the seasonal dynamics of *Ae. albopictus* and *Ae. aegypti* have been studied on regional scales for *Ae. albopictus* and *Ae. aegypti* in earlier studies [75–81]. By improving habitat suitability maps from yearly to monthly temporal resolution, the intra-annual seasonality of the distributions of *Ae. albopictus* and *Ae. aegypti* is captured through the model on a global scale. Unlike process-based mechanistic models that incorporate the life cycle and dispersal of mosquitoes [81], our methods rely on finding the species-specific niche for spatio-temporal mapping of *Ae. albopictus* and *Ae. aegypti* and fall in the category of correlative environmental niche models [75, 76, 80, 82]. We therefore compare our results with recent studies that utilize the correlative environmental niche modelling approach. In recent period, the seasonality of *Ae. albopictus* has been studied regionally over Europe by [80] and over China by [33] while the seasonality of *Ae. Aegypti* over United States has been covered in the work of [78]. On comparing the results of [80] over Europe with Figure 3A, it is found that the aggregated number of habitat suitability months as predicted by the model corroborates the findings of [80]. Although they have reported the seasonality of *Ae. albopictus* for the year 2019, our results are mostly consistent with their predictions over most of Europe and differ only over southern parts of Spain and Portugal. We find year-round habitat suitability of *Ae. albopictus* over these parts of Spain and Portugal whereas [80] reported unsuitable habitat conditions for *Ae. albopictus* over these regions. The difference in prediction results is likely due to the use of a precipitation suitability mask incorporated by [80], which rendered these parts of Spain and Portugal as unsuitable habitats for *Ae. albopictus*. On comparing the contemporary maps of *Ae. albopictus* over China presented in [33] with the results shown in Figure 3A, it is found that our results mirror their predictions of year-round habitat suitability in southern parts of China and low habitat suitability over western parts of China. Additionally, the decline in the number of habitat suitable months for *Ae. albopictus* over China with increasing latitude as seen in Figure 3A also validate the findings of [33]. The seasonality of *Ae. Aegypti* over United States has been studied by [78] and the predictions of our model largely support their findings, which shows year-round habitat suitability of *Ae. aegypti* over Florida and southern Texas. Over the west and east coast of United States, our results in Figure 1B show habitat suitability of *Ae. aegypti* between April and October and an aggregate of 7-8 months in Figure 3A, which compliments the maps of [78] for these regions. In other parts of United States, we find that the model predicts an increased northward distribution of *Ae. aegypti* during the summer months as compared to the results of [78], which is most likely a result of expanding habitat suitability of *Ae. aegypti* over the past ten years.

In addition to constructing global maps, we analysed and interpreted the contribution of individual covariates to model predictions, enabling us to examine the interplay between climate and land use variables in shaping the model’s decision. Our findings indicate that climatic variables, particularly temperature and dew-point temperature, exert a major influence on habitat suitability outcome of both *Ae. aegypti* and *Ae. albopictus*. From literature, it is known that the range of temperature and dew-point temperature values place a limiting factor on the survivability of both *Aedes* species. The Climademic Suitability Model is able to capture this aspect of mosquito species biology, which is discernible from the SHAP plots in Figure 3B. The extreme feature values of both these climatic variables lead to negative SHAP outputs, thus indicating that the model has learnt the suitable range of temperature and dew-point temperature necessary for the survivability of both these *Aedes* mosquitoes. However, the correct range of temperature and dew-point temperature do not solely determine the suitability of a given area. For example, the higher value of land use category “Sparse / No Vegetation” strongly shifts the model’s prediction toward lower suitability, even when temperature and dew-point temperature fall within otherwise favourable ranges. Similar to the results of [70], we find that the effect of the urban LULC variable do not play a major role in the decision-making process of the model for both *Ae. albopictus* and *Ae. aegypti*. Although, in literature, urbanicity is known to be an important covariate for the habitat of both *Ae. albopictus* and *Ae. aegypti* [83, 84], it is feasible that the grid cell size employed in this study is too coarse to sufficiently capture the urban/non-urban variation in a meaningful way, thus resulting in lower feature importance for this covariate. Additional finer scale studies need to be conducted to assess the importance of urbanicity on the habitats of *Ae. albopictus* and *Ae. aegypti*.

The present study provides the most recent and detailed reconstruction of the global potential distributions of *Ae. albopictus* and *Ae. aegypti*. Together, the improved spatio-temporal resolution and mechanistic explainability of the Climademic Suitability Model translates predictive outputs into actionable public health intelligence needed to guide the development of arbovirus surveillance and vector control programs. Additionally, knowing the intra-annual seasonality of the *Aedes* species, and the factors influencing it, provides crucial evidence to policy-makers for the design and implementation of cost-efficient and effective strategies to mitigate expansion and health risks posed by these invasive mosquito species. A first-order use of these maps can be found in Fig. 4, where we showcase the relationship and spatial overlap between human population and persistence in habitat suitable months of both *Aedes* species.

Averaging the change in suitability across locations between the decade of 1975-1984 and 2015-2024, our results suggest that inhabited areas are, on average, more suitable for both vector species. According to our model, approximately 5 billion people currently live in areas that have become more suitable habitats for both mosquito species since the mid 1970s. These areas also experienced the largest population growth with an addition of around 2 billion people in these areas, as shown in SI: Figure 7. The biggest share of population exposed to increasing suitability is concentrated in Asia and Africa, partly attributed to local rapid urbanization and demographic expansion. In Africa, approximately 678 million people currently live in such areas, of which around 391 million (ca. 58% of its entire population) represent population growth over the past 4 decades. In Asia, about 3.3 billion people inhabit such areas, with roughly 1.1 billion (ca. 34%) reflecting population increases over the same period. In contrast, in the European continent where over 92% of the population lives in areas that have since becoming more suitable for both species, demographic growth has been limited.

As a result, the increase in population living in areas that have became propitious to these insects is relatively smaller: approximately 39 million for *Ae. aegypti* and 33 million for *Ae. albopictus*. It is also worth noticing that our results point that currently around 34% of the world population, roughly 2.7 billion people, live in areas our model classifies as suitable for these insects all year around. As it can be seen in SI: Figure 8, this category has experienced the highest increase, both in absolute population and relative population growth.

In this work, both the yearly and monthly global maps were generated with the limitation that the LULC dataset was only available up to and including the year 2019. Consequently, we assumed that land use and land cover conditions changed sufficiently slowly to justify approximating the years 2020–2024 using the year 2019 LULC data. Another limitation arises from the model’s design: when estimating the suitability of a given 0.25° x 0.25° grid cell, the model does not incorporate information from adjacent cells. Considering the relatively limited flight range of these mosquito species, this assumption reflects the idea that even if neighbouring areas exhibit higher suitability, such conditions do not directly influence the species’ presence within the focal cell. A further constraint of our approach is that the model’s monthly suitability estimates do not account for suitability conditions in preceding months. This shortcoming might introduce inconsistencies, as mosquito activity is not governed by isolated, month-specific conditions. For instance, conditions in March may appear highly suitable, but exceptionally cold temperatures in January and February could markedly reduce overwintering egg survival or delay post-diapause development, preventing larvae from reaching adulthood in time to utilise favourable conditions. Such temporal dependencies may therefore contribute to discrepancies between predicted suitability and observed mosquito presence in validation datasets.

It is important to note that habitat suitability does not equate to actual mosquito presence. Rather, it represents environmental conditions for mosquito viability upon introduction. The models developed in this study were trained on mosquito sightings data from multiple sources that do not distinguish between established populations and transient introductions, therefore, over-prediction is possible. Furthermore, the use of quantile binning to transform the binary outputs of OCSVM models into calibrated probabilities comes with its own set of limitations. With the use of *B* bins, the probabilities are inherently limited to only *B* possibilities and there is an abrupt change in estimated probabilities as well at the boundaries of bins [85]. As we choose to use 10 bins for the quantile binning approach in this study, there is an intrinsic uncertainty of 0.1 in calibrated probability values, which must be taken into account when interpreting the results. Additionally, mosquito observations are inherently human-reported, and therefore spatially biased toward populated or urbanised areas. This bias has the potential to influence model performance. However, our SHAP analysis does not indicate that population density meaningfully contributes to model decisions. Nevertheless, this limitation should be taken into account when interpreting the results.

## 6 Summary and Outlook

We introduced the Climademic Suitability Model, a machine learning model that delivers monthly global predictions of *Aedes* habitat suitability at 0.25° spatial resolution between 1975—2024. Detailed analyses indicate that not only the majority of the world’s human settlements are located in areas that are becoming increasingly propitious for *Aedes* mosquitoes but also that the majority of the world population growth is happening in these areas, possibly exacerbating the contact between these arthropods and humans and future infectious disease burden to public health. Our findings suggest that the threshold of five billion people at risk of arboviral disease as projected to be reached by 2050 [86], may already have been surpassed. This finding is in line with concurrent global suitability estimates [87] and corroborating the urgency for coordinated vector control at scale.

Previous studies have used ecological niche modeling to constrain the spatiotemporal distribution of vectors for the development of virus-specific transmission models. The high-resolution spatial and seasonal information on the environmental suitability of the *Aedes* habitats gathered from this study can be applied to various downstream tasks and research directions, such as transmission studies for dengue, chikungunya and Zika viruses. Further developments to the suitability model are planned, including the addition of further climate covariate for improved estimates of habitat suitability of *Ae. albopictus* and *Ae. aegypti*, as well as the projected spatio-temporal distributions of these vectors for various climate scenarios.

## Data availability

All data used in this study is publicly available. All data used to train the Climademic Suitability Model are described in the data sections and references therein. The monthly suitability data fields of the model are available in a permanent data repository[88] via https://zenodo.org/records/19615975.

## Acknowledgements

The Federal Ministry of Research, Technology and Space (BMFTR) supports this study by funding the Climademic project (funding code 01LN2210A) within the framework of the Strategy Research for Sustainability (FONA). The Germany Federal Ministry of Health (BMG) supports this study by funding the AI-DAVis-PANDEMICS project (funding code 2523DAT400).

## Notes

https://zenodo.org/records/19615975

### Competing Interest Statement

The authors have declared no competing interest.

