## Supplementary Information for "Suitable seasons: Global monthly habitat suitability for the arbovirus vectors *Aedes aegypti* and *Aedes* albopictus in 1975–2024"

#### Supplementary Results

##### Model predictions: Annual habitat suitability

For the year 2024, the OCSVM yearly model predicts that the tropical and sub-tropical regions are broadly suitable for both *Ae. aegypti* and *Ae. albopictus* as shown in SI: Figure 1. These regions include equatorial and temperate parts of Africa and South America, Indian subcontinent and south east Asia, central America, mediterranean Europe and temperate parts of North America. The distributions of the two species are broadly similar for many regions all over the world but also differ markedly for a number of locations. The suitability of *Ae. aegypti* and *Ae. albopictus* differ mainly in Europe and Australia. While *Ae. aegypti* is found to be more suitable than *Ae. albopictus* in coastal parts of Eastern and Western Australia, in Europe, the distribution of *Ae. albopictus* extends more into higher latitudes in comparison to the distribution of *Ae. aegypti*. Such extended distribution of *Ae. albopictus* in comparison to *Ae. aegypti* is also seen in northern regions of United States, Japan and China.

With the mapping to probability levels, the predicted distributions of *Ae. aegypti* and *Ae. albopictus*, shown in SI: Figure 2, provide a better representation of the location of the specific hotspots of these two species. For *Ae. aegypti* and *Ae. albopictus*, these locations are concentrated in Eastern Brazil, central America, central Africa, central and Southern India, Southern China and coastal regions of South east Asia. The major differences in the distributions of these two species are seen in Europe, Eastern Australia and United States. While the greater suitability of *Ae. albopictus* over Europe than *Ae. aegypti* was also predicted in the binary classification plot, the probability levels clarify that only Mediterranean regions and parts of eastern Europe offer conditions that are moderately suitable for *Ae. aegypti*. On the other hand, the distribution of *Ae. albopictus* is predicted to cover much of Europe with hotspots in Mediterranean regions, southern France and central Europe. Over United States, *Ae. albopictus* is predicted to attain higher probability levels than *Ae. aegypti* over the eastern and central parts, while over Australia, *Ae. aegypti*, mostly confined to the north-east, is predicted to have higher probability levels than *Ae. albopictus*. By mapping the binary classification outputs of OCSVM yearly model to probability levels, the distributions of *Ae. aegypti* and *Ae. albopictus* provide a representation that is comparable to the global maps that are presented in Kraemer et al., (2015) [1].

### Supplementary Figures

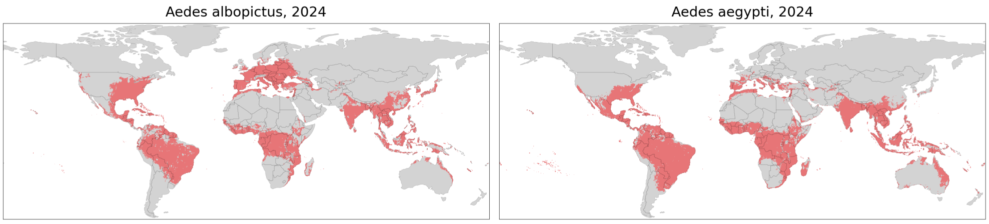


**Fig. 1**: For the validation year 2024, the binary classification outputs of the OCSVM yearly model are shown for *Ae. albopictus* and *Ae. aegypti* for all the global locations at a horizontal resolution of $0.25° \times 0.25°$ in this figure. The inlier points classified by the OCSVM yearly model are shown in red while the outlier points are shown in grey. The inlier points denote the locations that are predicted to be habitat suitable while the outlier points denote the locations that are unsuitable for *Ae. albopictus* and *Ae. aegypti*.

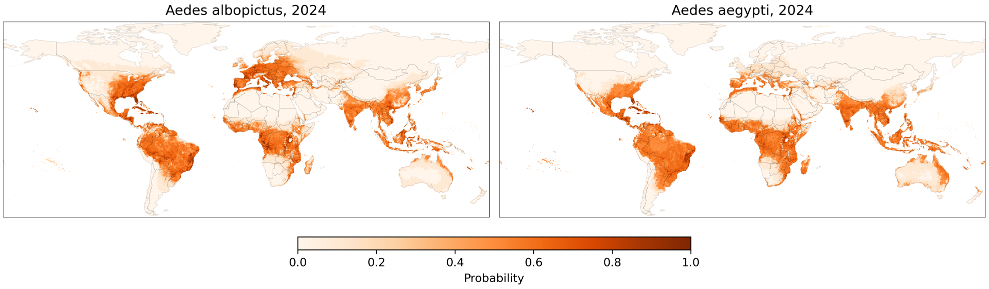


**Fig. 2**: For the validation year 2024, the binary classification outputs of OCSVM yearly model for *Ae. albopictus* and *Ae. aegypti* are mapped to probabilities and shown for all the global locations at a horizontal resolution of $0.25° \times 0.25°$ in this figure.

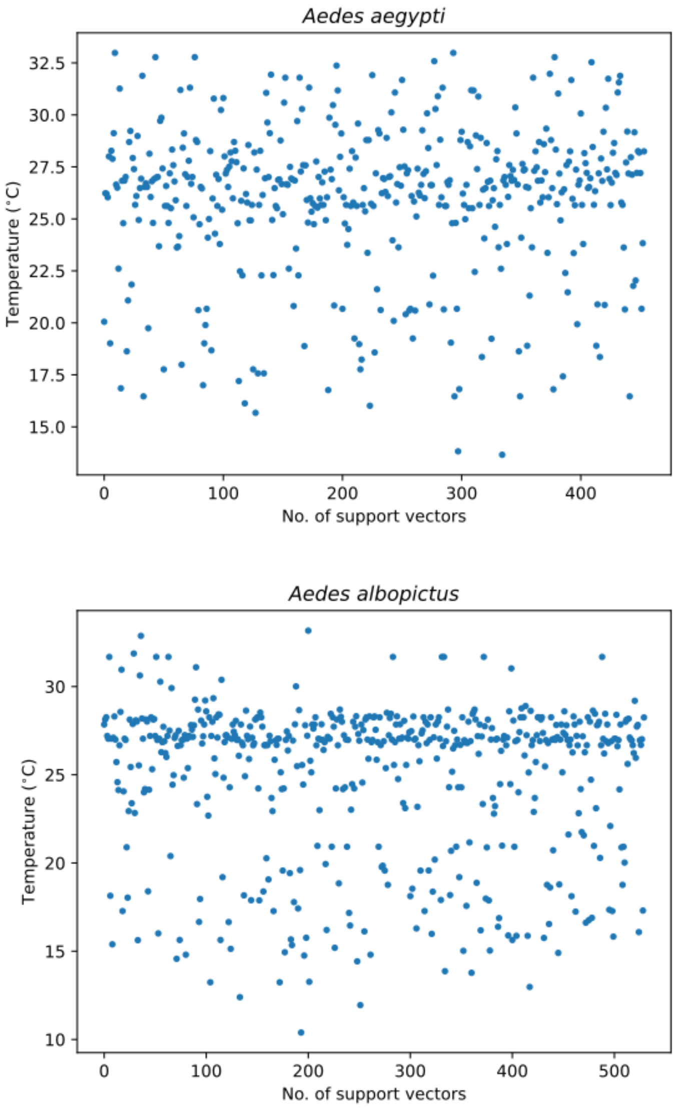


**Fig. 3**: Temperature range of support vectors with the choice of 75th percentile value for *Ae. aegypti* in the top panel and for *Ae. albopictus* in the bottom panel.

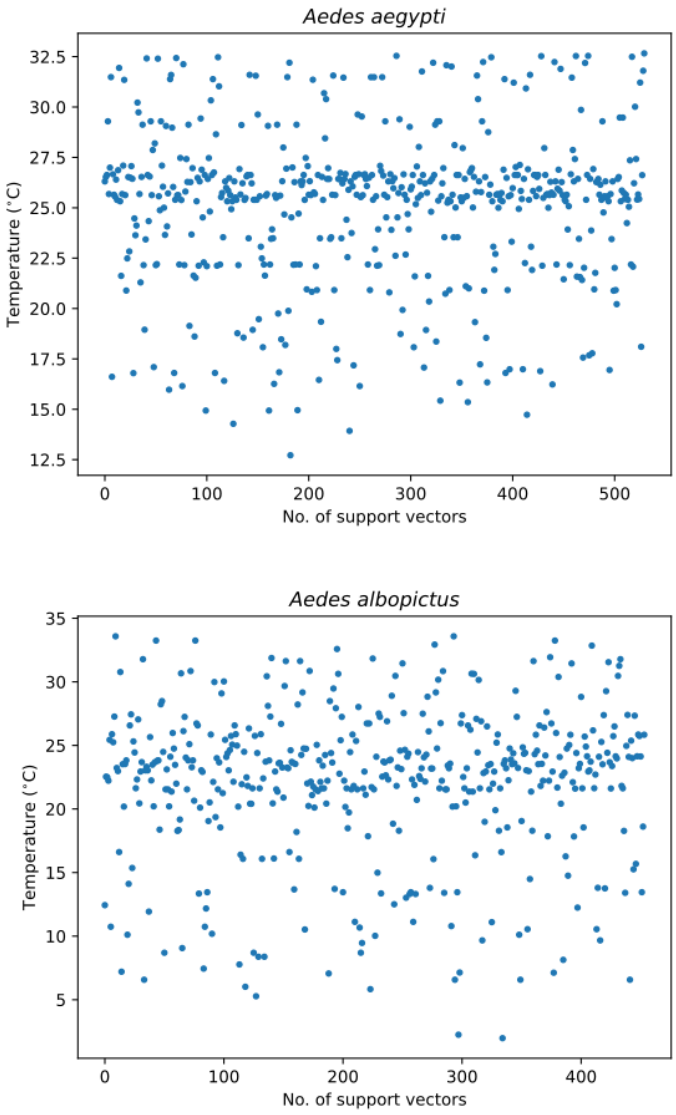


**Fig. 4**: Temperature range of support vectors with the choice of 50th percentile value for *Ae. aegypti* in the top panel and for *Ae. albopictus* in the bottom panel.

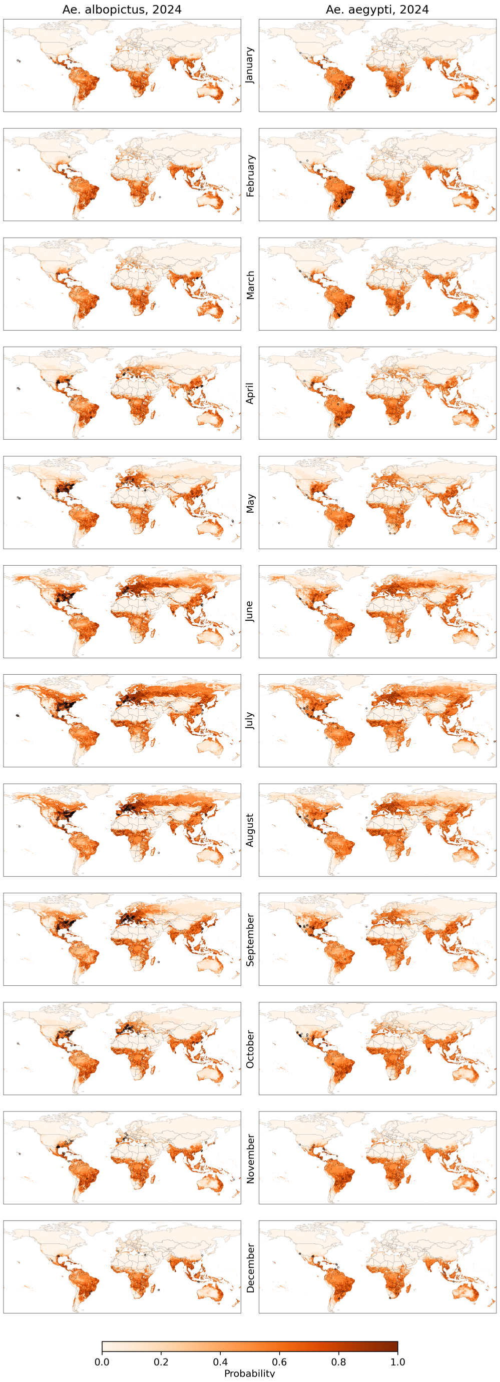


**Fig. 5**: Monthly habitat suitability for *Ae. albopictus* (left) and *Ae. aegypti* (right) for the months of January to December for the year 2024. Maps show posterior probability values of the OCSVM monthly model. Black rings indicate mosquito sightings, data based on X.

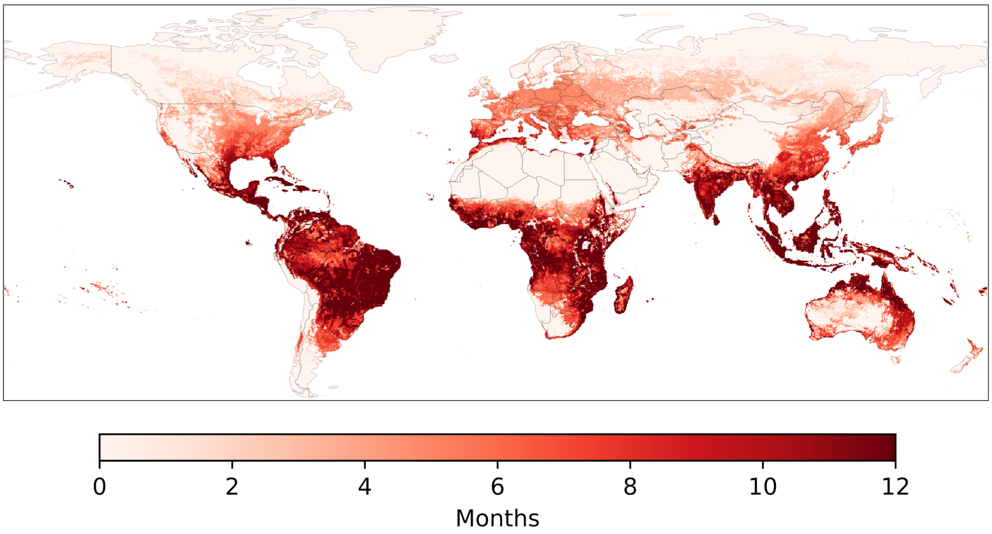


**Fig. 6**: Number of habitat suitable months at all global locations for *Ae. aegypti* for the year 2024.

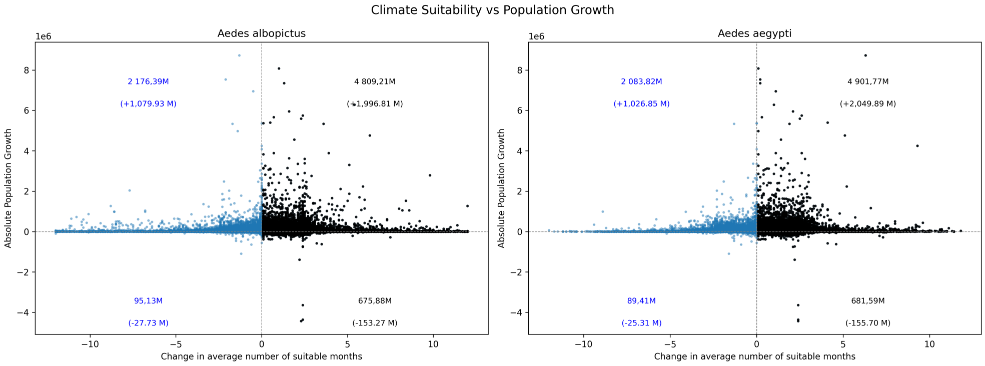


**Fig. 7**: Change in climatic suitability versus population growth across all grid cells for *Ae. aegypti* and *Ae. albopictus*. X-axis shows the change in the number of suitable months (between 1975–1984 and 2015–2024), and the Y-axis shows absolute population growth.

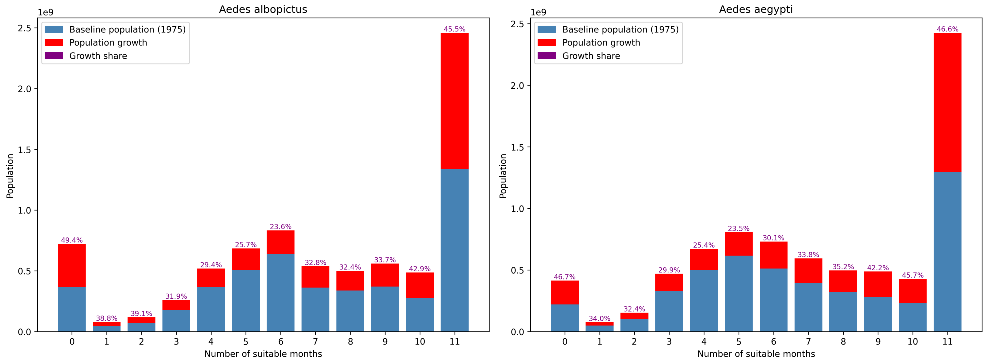


**Fig. 8**: Population distribution binned by the model estimated number of suitable months for *Ae. aegypti* and *Ae. albopictus*. Bars show baseline population in 1975 (blue) and subsequent population growth (red), stacked within each suitability bin. Purple labels indicate the percentage share of growth relative to total population in each bin since baseline population.
